# MEG State Dynamics of Sentence Generation: Evidence for a Compensatory Chunking Mechanism in Healthy Aging

**DOI:** 10.64898/2026.03.08.710384

**Authors:** Clément Guichet, Sylvain Harquel, Raouf Zouglech, Camille Lemaire, Émilie Cousin, Vincent Auboiroux, Aurélie Campagne, Monica Baciu

**Author notes:** **Corresponding Author:** Pr. Monica BACIU MD PhD, LPNC CNRS UMR 5105 Université Grenoble Alpes, 1251 rue des Universités, CS 40700, 38058 Grenoble Cedex 9 France.

## Abstract

Healthy aging is accompanied by subtle difficulties in language production. While behavioral and neuroimaging studies suggest that older adults rely on acute semantic access to maintain language abilities, the underlying neurophysiological mechanisms remain poorly understood. In particular, it is still unclear how large-scale brain dynamics reorganize to support naturalistic sentence generation with age. In this study, we investigated the spatiotemporal brain-state dynamics during covert sentence generation (GE2REC protocol) in younger and older adults using magnetoencephalography (MEG). Source-reconstructed MEG signals were analyzed using a Hidden Markov Model which identified five recurrent brain states, encompassing language-semantic, language-control, sensorimotor, and visual domains. Latent modeling was then used to relate the spectral and temporal properties of these brain states to age and language performance. Spectrally, older adults appear to redistribute oscillatory activity from sensorimotor-related states toward semantic-related states across alpha, beta, and low-gamma frequency bands. Temporally, older adults exhibit a more segmented processing sequence between semantic and sensorimotor processing which interfaces with visuo-posterior processing. These changes robustly covaried with age and better verbal fluency (semantic & lexical). Taken together, these results suggest that the older adult brain undergoes a coordinated time-frequency reorganization to support sentence production. Individuals likely establish an embodied semantic strategy in aging that involves “chunking” the processing stages of sentence production via visuo-posterior information processing. We speculate that this may help shape a resource-efficient, predictive route for complex cognition in older adulthood.

## 1. Introduction

The projected doubling of the world’s aging population by 2050 constitutes a major public health challenge, and consequently requires identifying the determinants of a healthy aging mind to inform preventive strategies (United Nations, 2023; WHO, 2025). Despite extensive behavioral and neuroimaging evidence, the neurophysiological mechanisms through which the aging brain dynamically supports complex cognition remain poorly understood.

Language production, as a central element of our neurocognitive system, offers a powerful framework to address this gap (Borne et al., 2024; Cui et al., 2025; Hagoort, 2023). At the behavioral level, language is increasingly used as a biomarker of the strengths and vulnerabilities in healthy and pathological aging (Benítez-Burraco & Ivanova, 2024; Mekkes et al., 2024). At the cognitive level, language involves cross-domain interactions between domain-general executive functioning, domain-specific semantic processing, and perceptuo-motor systems (Bayram et al., 2023; Bourguignon & Lo Bue, 2025; Roger et al., 2022). Therefore, language is an ideal cognitive function to investigate interacting processes, especially those drawing from domain-general and domain-specific systems in aging, such as cognitive flexibility (Endress, 2019; Guichet et al., 2025). This cross-domain architecture is further evidenced at the brain level: the classical view of a unitary “language network” is increasingly being replaced by a connectomic, whole-brain anatomo-functional description (Aliko et al., 2023; Forkel & Hagoort, 2024; Hagoort, 2019; Thiebaut de Schotten & Forkel, 2022). Accordingly, the present study focuses specifically on language production as an indicator of how aging reshapes the dynamic coordination between semantic, control, sensorimotor and perceptual systems throughout brain.

Language production is particularly vulnerable to aging (Baciu et al., 2021). Behaviorally, these difficulties manifest as increased picture-naming latencies and a higher incidence of tip-of-the-tongue states beyond midlife (Benítez-Burraco & Ivanova, 2023), especially for proper names and semantically isolated words (Bannon et al., 2024; Brown & McNeill, 1966). Sentence production is similarly affected (Peelle, 2019), especially for syntactically complex structures and tasks with high working-memory demands (Agmon et al., 2023; Kemper et al., 2004; Sung, 2015). A leading hypothesis attributes these difficulties to age-related decline in executive control (Facal et al., 2012; Rossi & Diaz, 2016), which compromise the inhibition of competing representations during lexico-semantic access (Baciu et al., 2016; Badre & Wagner, 2007; Blanco et al., 2016; Boudiaf et al., 2018; Hoffman & Morcom, 2018). For example, older adults have more difficulty resolving co-activated lexical competitors and are more susceptible to semantic interference (Federmeier et al., 2020; Hardy et al., 2022; Sung, 2015). Critically, converging evidence shows that these age-related difficulties reflect a reorganization of the underlying brain dynamics that support lexical and sentence-level production rather than a uniform loss of neural efficiency.

Notably, a large body of work suggests that the lifelong accumulation of semantic knowledge may compensate some of the language production difficulties (Boudiaf et al., 2018; Cutler et al., 2025; Gilis et al., 2025; Gollan & Goldrick, 2019; Krethlow et al., 2024). This “semantic strategy” for language production can be described at complementary levels: (i) behaviorally, Fargier & Laganaro (2023) demonstrated that older adults perform better when the word production is driven by semantic associations (i.e., inferential production) than in traditional picture naming tasks (i.e., referential production); (ii) at the brain level, the LARA-C model (Baciu et al., 2021; Baciu & Roger, 2024) highlights the recruitment of age-resilient ventral semantic pathways, supporting top-down modulation from inferior frontal to medial temporal regions to facilitate semantic memory access (Hoyau et al., 2018). Within a connectomic framework, the SENECA model offers a complementary framework that emphasizes how semantic processing also relies on embodied and perceptual systems with age (Guichet et al., 2024, 2026), echoing earlier evidence showing that sensory-perceptual enrichment can partially offset control decline in older adults (Baltes & Lindenberger, 1997; Lindenberger & Baltes, 1994; Lindenberger & Mayr, 2014). Taken together, behavioral, cognitive, and neuroimaging findings support that age-related language production reflects an adaptive reorganization rather than a uniform deterioration of our brain. Age-related decline in executive control coexists with the recruitment of semantic and sensorimotor pathways for preserving lexical access and retrieval (Guichet et al., 2025b).

From a neurophysiological perspective, understanding how aging reshapes language production requires methods capable of capturing fast, transient and large-scale brain dynamics. Magnetoencephalography (MEG) is particularly suited for this purpose, as it allows the characterization of oscillatory regimes during language tasks. Relatedly, recent MEG work suggests that age-related semantic processing is grounded in predictive processing. For example, a resting-state MEG study showed that acute semantic access in older adulthood correlates with increased beta to low-gamma band dynamics (13-45 Hz) across all brain states (Guichet et al., 2025). This result is consistent with prior work linking beta-gamma synchrony to post-stimulus prediction error (Fujioka et al., 2009), and implicating gamma-band oscillations in sensory prediction errors and perceptual integration (Bastos et al., 2012; Chao et al., 2022; Jensen et al., 2014). In addition, a task-MEG study reported decreases in left temporal-parietal beta power in older adults (Zheng & Piai, 2025). These beta suppressions were interpreted as a semantic integration mechanism (Hanslmayr et al., 2009; Piai & Zheng, 2019), potentially reflecting the reactivation of content representations during syntactic and semantic context integration (Antzoulatos & Miller, 2016; Spitzer & Haegens, 2017; Zioga et al., 2023).

In sum, beta-gamma-range modulations appear as crucial substrates for establishing the hypothesized semantic strategy in older adults: beta desynchronization may facilitate the propagation of sensory prediction for semantic integration through gamma-mediated signaling (Bornkessel-Schlesewsky & Schlesewsky, 2019; Wei et al., 2025). However, the MEG evidence for this semantic strategy remains inconclusive, whether based on resting-state (Guichet et al., 2025) or picture-word interference (Zheng & Piai, 2025). This highlights the need to investigate the neural dynamics of language production in a more naturalistic fashion, embedding lexical retrieval within a continuous semantic and syntactic context.

To specify the neurophysiological mechanisms supporting the semantic strategy in aging, we adapted a naturalistic sentence generation paradigm initially performed in fMRI (GE2REC protocol; Banjac et al., 2021) to MEG. Moreover, we modeled MEG activity using a latent whole-brain dynamics approach that integrates spatio-spectral and temporal properties of the data in an unsupervised fashion. We hypothesized that sentence generation engages multiple recurrent brain states whose properties differ between younger and older adults: spectrally, we expected age-related modulations in the beta and low-gamma oscillatory regimes, correlating with a semantic strategy for sentence generation. Temporally, we expected a distinct cyclical architecture between brain states with age, reflecting how the processing stages of sentences production are re-organized to establish the semantic strategy.

## 2. Material and Methods

### 2.1. Participants

MEG recordings were acquired from 22 healthy participants divided into a younger adult group (*N* = 12, 6 females, 6 males, mean age = 28.3 yo, *SD* = 4.2) and an older group (*N* = 10, 4 females, 6 males, mean age = 65.2 yo, *SD* = 6). All participants were native French speakers, had normal or corrected-to-normal vision, and were right-handed (laterality was assessed using Edinburgh Handedness Inventory; Oldfield, 1971). All participants provided written informed consent prior to participation. The study was approved by the local ethics committee of CHU Grenoble Alpes (CPP MEG-AGING n° *38RC18.294*, CHUGA).

Participants had no history of neurological disease, no major psychiatric disorders, and met a cognitive inclusion criterion defined by a Mini-Mental State Examination (MMSE) score greater than 25/30 (Kalafat et al., 2003). Anxiety and depressive symptoms were assessed using the HAD scale (Snaith & Zigmond, 1986) and added as covariates in statistical models. Cognitive abilities were evaluated using standardized measures for language, including vocabulary with the Mill-Hill test (Deltour, 1993), lexical and semantic fluency (Cardebat et al., 1990), and executive functions with the BREF (Dartinet & Martinaud, 2005) and the TMT (Tombaugh, 2004) (see Table 1).

**Table 1.**
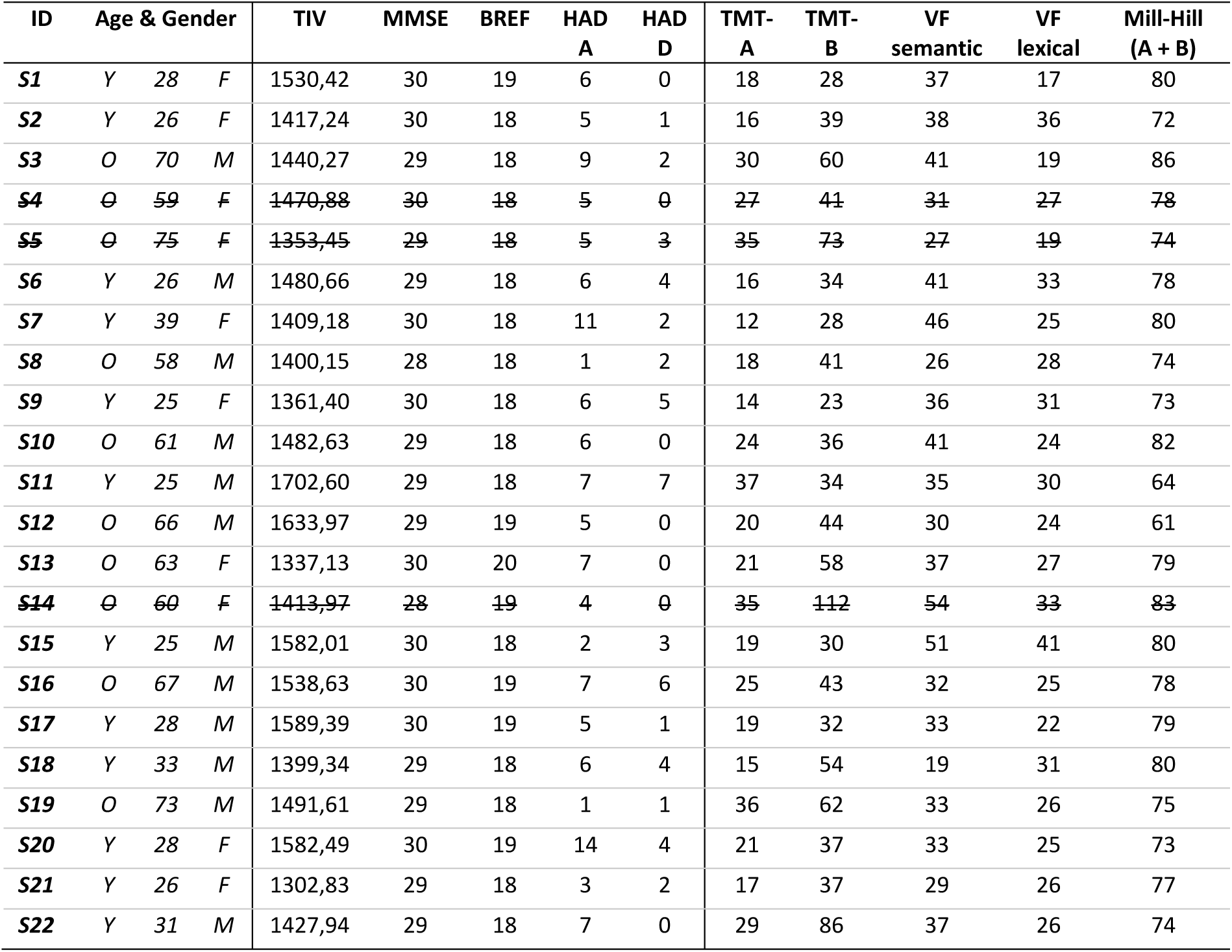
Demographic and Neuropsychological characteristics of the participants. Barred subjects (S4, S5, S14) were excluded for statistical analysis due to strong aperiodic activity compared to the other subjects (see section 2.5). *Y: Younger; O: Older; F: Female; M: Male; TIV: Total Intracranial Volume; VF: Verbal Fluency*

| ID | Age & Gender |  |  | TIV | MMSE | BREF | HAD<br>A | HAD<br>D | TMT-A | TMT-B | VF<br>semantic | VF<br>lexical | Mill-Hill<br>(A + B) |
| --- | --- | --- | --- | --- | --- | --- | --- | --- | --- | --- | --- | --- | --- |
| S1 | Y | 28 | F | 1530,42 | 30 | 19 | 6 | 0 | 18 | 28 | 37 | 17 | 80 |
| S2 | Y | 26 | F | 1417,24 | 30 | 18 | 5 | 1 | 16 | 39 | 38 | 36 | 72 |
| S3 | O | 70 | M | 1440,27 | 29 | 18 | 9 | 2 | 30 | 60 | 41 | 19 | 86 |
| ~~S4~~ | ~~O~~ | ~~59~~ | ~~F~~ | ~~1470,88~~ | ~~30~~ | ~~18~~ | ~~5~~ | ~~0~~ | ~~27~~ | ~~41~~ | ~~31~~ | ~~27~~ | ~~78~~ |
| ~~S5~~ | ~~O~~ | ~~75~~ | ~~F~~ | ~~1353,45~~ | ~~29~~ | ~~18~~ | ~~5~~ | ~~3~~ | ~~35~~ | ~~73~~ | ~~27~~ | ~~19~~ | ~~74~~ |
| S6 | Y | 26 | M | 1480,66 | 29 | 18 | 6 | 4 | 16 | 34 | 41 | 33 | 78 |
| S7 | Y | 39 | F | 1409,18 | 30 | 18 | 11 | 2 | 12 | 28 | 46 | 25 | 80 |
| S8 | O | 58 | M | 1400,15 | 28 | 18 | 1 | 2 | 18 | 41 | 26 | 28 | 74 |
| S9 | Y | 25 | F | 1361,40 | 30 | 18 | 6 | 5 | 14 | 23 | 36 | 31 | 73 |
| S10 | O | 61 | M | 1482,63 | 29 | 18 | 6 | 0 | 24 | 36 | 41 | 24 | 82 |
| S11 | Y | 25 | M | 1702,60 | 29 | 18 | 7 | 7 | 37 | 34 | 35 | 30 | 64 |
| S12 | O | 66 | M | 1633,97 | 29 | 19 | 5 | 0 | 20 | 44 | 30 | 24 | 61 |
| S13 | O | 63 | F | 1337,13 | 30 | 20 | 7 | 0 | 21 | 58 | 37 | 27 | 79 |
| ~~S14~~ | ~~O~~ | ~~60~~ | ~~F~~ | ~~1413,97~~ | ~~28~~ | ~~19~~ | ~~4~~ | ~~0~~ | ~~35~~ | ~~112~~ | ~~54~~ | ~~33~~ | ~~83~~ |
| S15 | Y | 25 | M | 1582,01 | 30 | 18 | 2 | 3 | 19 | 30 | 51 | 41 | 80 |
| S16 | O | 67 | M | 1538,63 | 30 | 19 | 7 | 6 | 25 | 43 | 32 | 25 | 78 |
| S17 | Y | 28 | M | 1589,39 | 30 | 19 | 5 | 1 | 19 | 32 | 33 | 22 | 79 |
| S18 | Y | 33 | M | 1399,34 | 29 | 18 | 6 | 4 | 15 | 54 | 19 | 31 | 80 |
| S19 | O | 73 | M | 1491,61 | 29 | 18 | 1 | 1 | 36 | 62 | 33 | 26 | 75 |
| S20 | Y | 28 | F | 1582,49 | 30 | 19 | 14 | 4 | 21 | 37 | 33 | 25 | 73 |
| S21 | Y | 26 | F | 1302,83 | 29 | 18 | 3 | 2 | 17 | 37 | 29 | 26 | 77 |
| S22 | Y | 31 | M | 1427,94 | 29 | 18 | 7 | 0 | 29 | 86 | 37 | 26 | 74 |

### 2.2. Experimental Protocol

The GE2REC (Generation, Recognition, Recall) protocol, initially developed for fMRI (see Banjac et al., 2021 for details) was adapted to MEG to investigate the neural dynamics of the language-memory interaction. We focused our study on analyzing the process of “Sentence Generation” because it simultaneously involves lexical retrieval, semantic integration, and syntactic planning, making it particularly well suited for investigating the neural dynamics of language in a naturalistic fashion.

Figure 1 illustrates the block design, which consisted of 40 randomized trials of 9 s for a total of 6 min per block. A block alternated between the task (20 trials with words) and control (20 trials with pseudowords) conditions. The entire protocol had 4 blocks with a resting period in between during which participants fixated a cross on a black screen. During the task condition, participants covertly generated short sentences for 5s (from *t* = 3 s to *t* = 8 s) after hearing a word at *t* = 0 s. Covert sentence generation allowed us to efficiently probe sensorimotor coordination, minimizing the articulatory-motor artifacts and external feedback while still preserving a temporal structure equivalent to the one observed in overt speech (Mantegna et al., 2026). Words were taken from the French standardized naming test DO80 (Deloche & Hannequin, 1997). During the control condition, participants listened to and repeated a pseudoword (/*mis-toudin*/) covertly instead of generating sentences.

**Figure 1.**
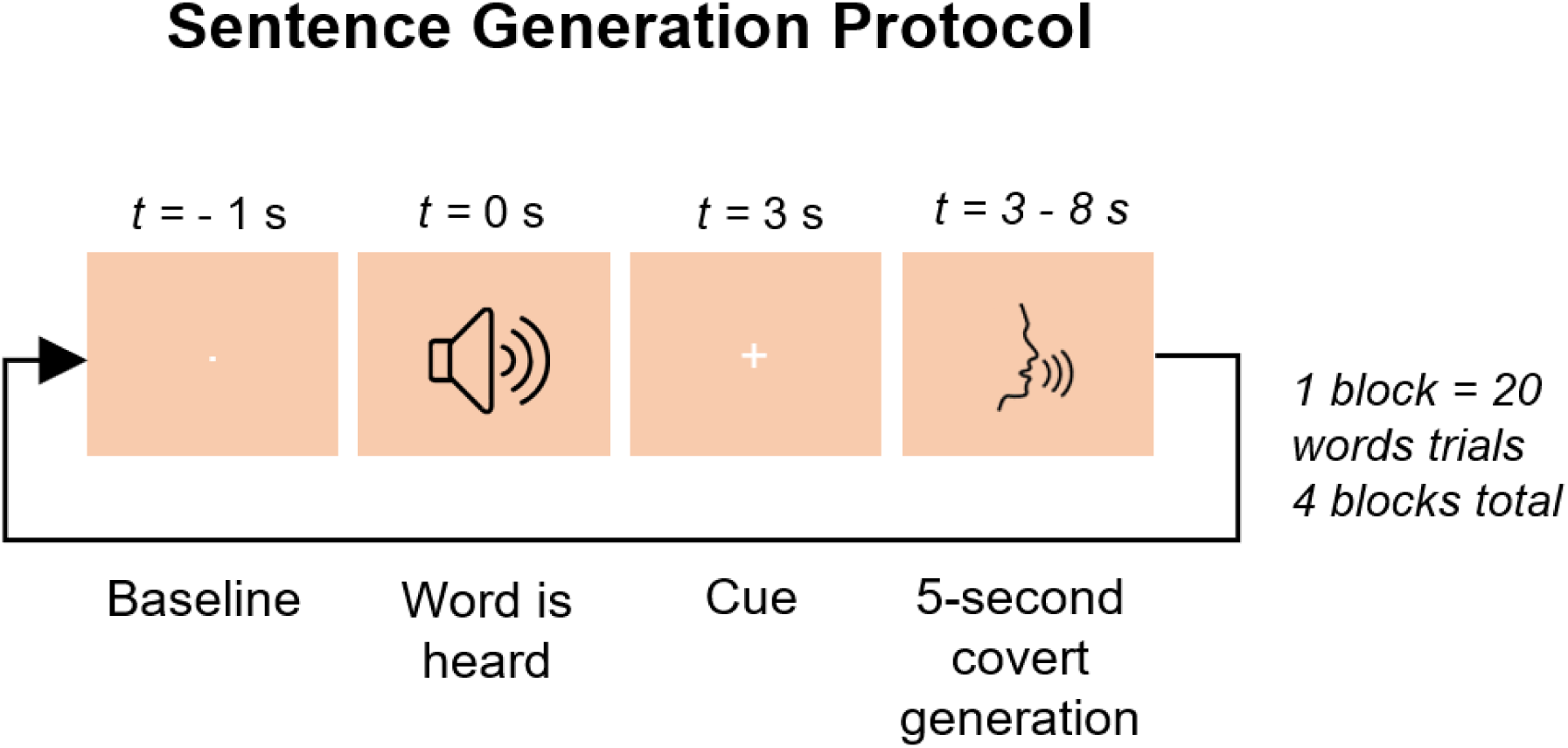
Schematic of the experimental protocol. Stimuli (cue words) were presented in the auditory modality, then sentences using the cue word were covertly generated for 5 seconds. We focused on the covert sentence generation period in this study.

### 2.3. MEG data acquisition and processing

#### Acquisition

MEG data were collected in magnetically shielded room in a 306-channel Vector-view MEG system (Elekta Neuromag), 102 magnetometers, and 204 orthogonal planar gradiometers (Clinatec Neuroimaging Facility). Data were sampled at 1 kHz.

#### Preprocessing

Preprocessing steps were initially carried out with the Brainstorm software (http://neuroimage.usc.edu/brainstorm), in MATLAB (Tadel et al., 2011). Temporal signal space separation (tSSS) (Taulu & Simola, 2006) was applied for noise reduction and offline head motion correction based on four head position indicators (HPIs). Data were notch filtered at 50 Hz and harmonics to remove power line noise. Further denoising was performed with Infomax ICA decomposing each block signal into 50 components before rejecting ECG/EOG-related artifacts.

After the removal of bad trials (visualization), remaining good trials were further preprocessed with MNE python. First, we applied bandpass filtering using a 5^th^-order IIR Butterworth filter [0.5 – 125 Hz] and downsampled to 250 Hz. Second, bad channels were interpolated using spherical spline interpolation (Perrin et al., 1989) to keep data dimensions consistent across participants. Third, each preprocessed trial was cropped between *t* = 3.5 and *t* = 8 s, thereby omitting the first 500 ms of the task condition. This was done to isolate the core sentence generation phase for downstream analyses, while excluding early evoked visuo-sensory responses common across participants related to the display of the cue (Fig. 1). Finally, all trials were concatenated along the time dimension to obtain one continuous trial sequence for each participant.

#### Co-registration

T1-weighted MR images were acquired with Siemens Espree 1.5T MR imager during a 30-min scan following the experimental procedure: TR=14 ms, TE = 4.87 ms, FA=25°, spatial resolution=1 mm isometric, FOV= 256, number of slices=176. MEG-T1 co-registration was conducted using digitized anatomical fiducial points (bilateral pre-auricular points, nasion, headshape) by extracting the scalp’s surfaces, inner skull, and brain with FSL Brain Extraction Tool (Smith, 2002). We used a 2-mm T1w MNI template for 3 participants (2 older, 1 younger) due to bad skull stripping and manually validated all co-registration outputs.

#### Source reconstruction

Following OHBA Software Library (OSL) guidelines, preprocessed data were bandpass filtered [1–45 Hz], and sources were reconstructed onto an 8-mm isotropic dipole grid. This reconstruction was based on a single-shell lead-field model in MNI space and employed a linearly constrained minimum variance (LCMV) scalar beamformer (Van Veen et al., 1997; Woolrich et al., 2011). Voxels were parcellated into 52 regions derived from the HCP-MMP 1.0 atlas (Kohl et al., 2023) in line with previous team work on lexical production (Guichet et al., 2025). Symmetric multivariate leakage reduction was applied to mitigate source leakage (i.e., artefactual correlations between parcel time courses) (Colclough et al., 2015), and dipole sign ambiguity was resolved using a sign-flipping algorithm (Vidaurre et al., 2018).

### 2.4. MEG analysis: Hidden Markov Model

A Hidden Markov Model (HMM) was employed to model MEG activity. This model assumes that oscillatory activity can be generated from a sequence of distinct brain states whose activations vary over time. As recommended for task data, we selected the DyNeMo variant which allows states to overlap at each timepoint instead of being mutually exclusive, and explicitly accounts for long-range temporal dependencies.

For a detailed mathematical description of the observation model, please refer to Gohil et al. (2022). All modeling steps were carried out with the OHBA Software Library Dynamics Toolbox (*osl-dynamics*; Gohil et al., 2023).

#### Data preparation

We prepared source-space data by adding time-lagged versions of each channel. These lags allowed the inference of spectral properties such as oscillatory amplitude and phase synchronization. As recommended, we used 15 lags evenly distributed around each timepoint across the range −125 Hz to 125 Hz. Considering our 52 parcels, this introduced 728 (14×52) off-diagonal elements to the covariance matrix used for model training. To prevent overfitting, principal component analysis (PCA) reduced this space to twice the number of parcels (i.e., 104) before z-scoring the data across the time dimension.

#### Model inference

Model hyperparameters were fine-tuned across the first runs based on training stability and convergence and subsequently set as follow: *sequence length*=100, *batch size*=16, *epochs*=50, *learning rate*=1e-3. We conducted 10 runs for *S* = 4, 6, or 8 states to find the optimal number of brain states fitting our data. We selected *S* = 6 states as the best compromise between model complexity and neurofunction interpretability based on the observed spatio-spectral features. As expected with our small sample size, we observed substantial variability across the 10 runs. Therefore, as recommended by Alonso & Vidaurre (2023), we employed hierarchical clustering to derive a more reliable solution that considers all runs rather than choosing the single best one.

### 2.5. MEG post-hoc analysis

#### Power spectral density

Once trained, the model provides a timeseries of the mixing coefficients, indicating the mixing ratio between brain states at each time point (e.g., 20% State 1, 10% State 2, etc.). To extract the power spectrum density (PSD) of each state, we first computed the Welch spectrogram from source-space data and regressed it on the mixing coefficients using the GLM-spectrum approach (Quinn et al., 2024). The regression coefficients can then be interpreted as state-specific PSDs for each subject.

#### Outliers

Initial inspection showed that State 4 resumed high-amplitude aperiodic activity across the whole-brain, whose parcel-averaged power was 3.62 z-units away the mean power observed in the other five states. This aperiodic profile was further confirmed by visualizing the subject-specific PSDs (see Figures S1 & S2, Supplementary Information). Relatedly, we excluded three outliers in the older age group which showed strong aperiodic activity in this state compared to the other subjects (see Figure S2, Supplementary Information). This brings the final sample size to *N* = 7 in this age group. These exclusions were done prior to any statistical analysis linking the states to behavioral variables, avoiding any circular bias. Because the regression coefficients are estimated with respect to the mean of all states, we refitted the GLM-spectrum after excluding this aperiodic state to focus only on oscillatory activities. This brings the final number of states to *S* = 5.

#### Network composition

To describe the states in terms of brain networks, we extended a method validated in previous teamwork (Guichet et al., 2025). First, we computed subject-specific power maps for each state by averaging each PSD across frequencies. Second, we computed the volumetric overlap between our 52 parcels and a 9-resting-state-network (RSN) atlas (Ji et al., 2019). The network composition of each state and subject was obtained by calculating the inner product between each subject’s power map (*dim:* 5 states x 52 parcels) and the volumetric overlap (*dim:* 52 parcels x 5 networks). We then averaged across subjects to obtain a single group-level network description whereby each state is associated with many resting-state networks (*dim*: 5 states x 9 networks). This approach allowed us to explicitly take into account the inter-individual variability in the states’ spatial layout.

### 2.6. Statistics

To examine how age and cognitive performances jointly modulate task dynamics, we employed Partial Least Squares correlation analysis (https://github.com/MIPLabCH/myPLS) in MATLAB R2025b. PLS is an unsupervised multivariate technique that extracts latent components by maximizing the covariance between two datasets: brain features (*X matrix*) and a behavioral profile comprising age and cognitive scores (*Y matrix*). PLS offers a principled way to manage: (i) multicollinearity across features, (ii) contexts where there are more features than participants (Mihalik et al., 2022); and (iii) enhances the robustness of individual differences (DeYoung et al., 2025).

#### PLS inference

The covariance matrix (R = X*Y^T^) was decomposed using singular value decomposition (SVD), resulting in R = USV^T^. Each latent component is defined by a singular value (s), representing the shared covariance, and a set of brain (u) and behavioral (v) saliences that denote the contribution of each feature to the component. Statistical significance of each component was assessed using 10,000 permutations, and p-values were adjusted at False Discovery Rate (FDR) across the components of each PLS model. Robustness of saliences was evaluated using 1,000 bootstrap resamples. We then calculated a Bootstrap Sampling Ratio (BSR) for each feature, defined as the observed salience weight divided by its bootstrapped standard deviation. Features with a BSR ± 3 were considered robust contributors, corresponding to approximately a 99% confidence interval (Krishnan et al., 2011).

#### Behavioral matrix preparation

The behavioral matrix (Y) was constructed from performance scores on the TMT A/B, as well as Verbal and Semantic Fluency tests. TMT scores were inverted so that higher values consistently indicated better performance. To isolate the specific effect of age and cognition, we regressed several covariates: anxiety and depression levels (HADS), gender, total intracranial volume (TIV), MMSE, BREF score, and Mill-Hill vocabulary score. The resulting residuals were quantile-normalized to ensure Gaussianity. Finally, age was appended as a continuous variable and the entire matrix was z-scored across subjects.

#### Brain feature matrix

This behavioral dataset was examined in relation to both spectral and temporal brain state properties through two PLS models:

- (*Model 1*) Spectral model: For each subject and state, we concatenated the PSD profile and z-scored across subjects. We limited our analysis to the top 10 parcels to ensure spectral relevance, that is focusing on each state’s highest group-average activity.
- (*Model 2*) Temporal model: We developed a method to capture non-trivial dynamic patterns organized in cycles, beyond simple pairwise state-to-state transitions. This allowed us to examine how sentence generation processes are chunked in time.

First, we computed the inner product of the state time courses (α) to obtain the transition probability matrix, *T*. To focus exclusively on state-to-state dynamics, we zeroed the diagonal elements and then normalized each row to obtain a jump-transition matrix, *P*:

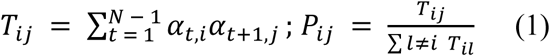

where *N* denotes the total number of timepoints in the session, and the denominator in the second expression represents the total flux out of state *i* into any other state *l*, excluding self-transitions.

This approach leverages the full posterior distribution of the HMM, modeling ‘soft’ transitions rather than the hard assignments typically used with binarized time courses, thereby respecting the model’s uncertainty. Considering the 5 states identified in this study (after exclusion of aperiodic activity), we generated a list of *M* = 60 unique cycles of lengths *k* ∈ {2,3,4}. For each cycle *C*, the probability ø(*C*) is calculated as the geometric mean of the transition probabilities along the sequence of states 〈*s*_*j*_, …, *s*_*k*_〉:

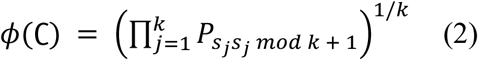

This formulation ensures the metric is normalized for cycle length and captures non-trivial dynamics as cycles containing ‘weak link’ transitions are penalized (if any link is near zero, the entire metric approaches zero). Because the resulting cycle values exist in a compositional space, we applied the centered-log-ratio (CLR) transform to linearize the data:

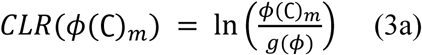

where *g*(ø) is the geometric mean of all computed cycles:

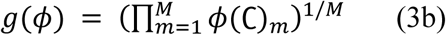

This procedure was repeated for each subject and the resulting cycle values were z-scored across subjects in preparation for PLS.

## 3. Results

### 3.1. Description of the brain states during sentence generation

The Hidden Markov Model revealed five recurrent brain states during the sentence generation period. Figure 2 illustrates the states’ network topography and spectral signature (see also Figs. S3 & S4, Supplementary Information).

**Figure 2.**
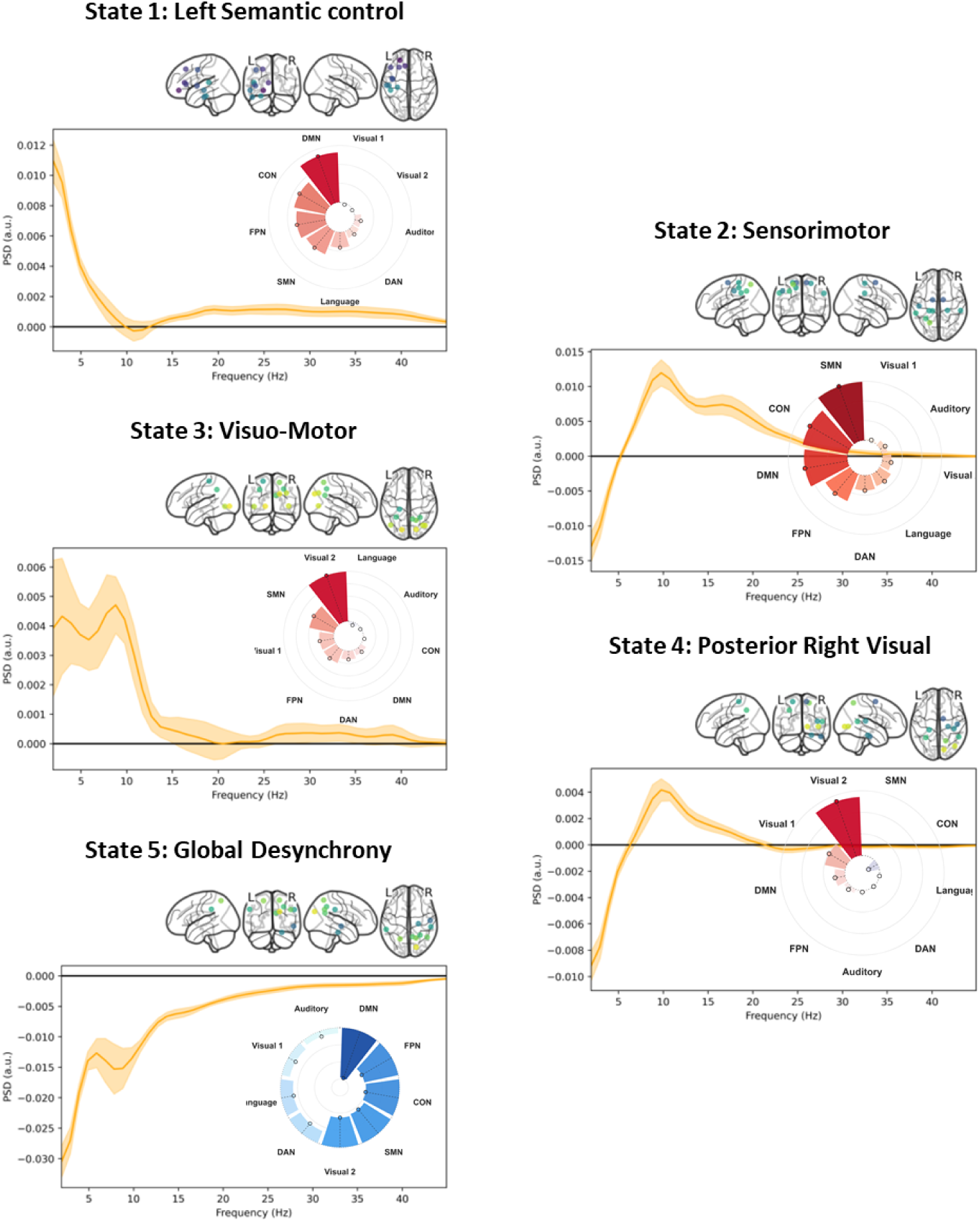
Dynamic spatio-spectral profiles of Sentence Generation. The main plot illustrates the group-average power spectra of each state across all 52 channels. Bandwidths represent the standard error across subjects. **Brain plot.** Illustrates each state’s 10 most-powered channels on average across all subjects. **Radar plot.** Illustrates the resting-state network topography of each state (red = synchrony, blue = desynchrony). *DMN (Default Mode), FPN (Fronto-Parietal), CON (Cingulo-opercular), DAN (Dorso-Attentional), SMN (Sensorimotor)*

State 1 highlights overlaps between left-lateralized communication between the DMN and attentional systems (FPN & CON) in low frequencies (delta/theta bands < 8 Hz) and aperiodic activity (1/f slope), a regime typically associated with top-down cognitive control and lexico-semantic retrieval (Cavanagh et al., 2012; Nigbur et al., 2011; Zheng & Piai, 2025). State 2 engages similar overlaps with the DMN and attentional systems (FPN & CON), but with the SMN as the main driver in a broad alpha-beta band (10-30 Hz).

States 3 & 4 are visuo-posterior states dominated by alpha activity with clear peaks in the 8-12 Hz band. State 3 specifically binds with the SMN with an additional peak in the delta range whereas State 4 binds with the DMN. Finally, State 5 represents a period of right-dominant desynchronization with respect to the mean task activation, which appears as the counterpart of DMN synchronization in State 1.

Initial cluster-permutation tests revealed no significant differences between age groups in neither of the five states (Fig. 3). In the next section, we employed a latent modeling approach to capture more holistic patterns of change across the state space, considering the contribution of all states simultaneously.

**Figure 3.**
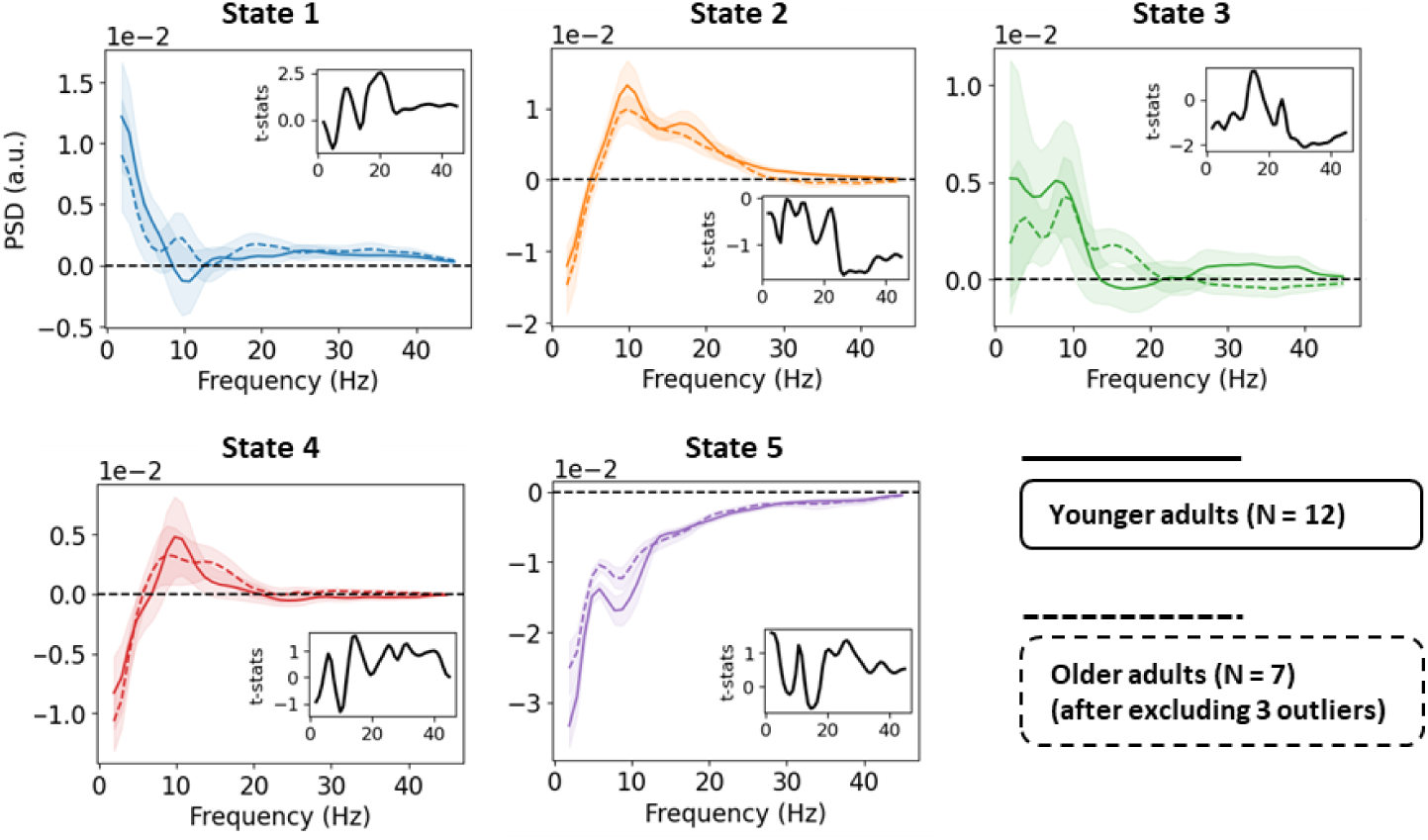
Cluster-based permutation results between age-group spectra. For each state, the plain line depicts the average spectrum of younger adults, and the dotted one that of older adults. The in-plot reports the t-statistic from 5,000-permutations. No significant clusters were identified following Bonferroni correction across states

### 3.2. Latent modeling between brain features, age, and cognitive performances

Together, the five states identified above constitute the basic dynamic units of the sentence generation task. Using the Partial Least Squares (PLS) analysis, we examined how their spectral properties and temporal organization covary with age and behavioral performances.

The PLS models identified a single significant latent component (*p_spectral_* = 0.016 & *p_temporal_* = .015), explaining respectively 39.5% and 50% of the total shared covariance. While neither component survived FDR correction (*pFDR_spectral_* = 0.079 & *pFDR_temporal_* = .075), spectral and temporal models converged on a similar behavioral profile. We begin by describing this behavioral profile, and then report the associated changes in oscillatory properties and sequential brain state organization.

#### 3.2.1. Behavioral profile

Overall, results suggest a coordinated time-frequency reorganization of sentence generation linked to age-related verbal fluency performances (Figure 4). Spectral features most robustly covaried with semantic and verbal fluency (BSR*_spectral_* = 17.23 & 9.25), whereas the temporal dynamics between states more robustly captured the effects of brain aging (BSR*_temporal_* = 5.75).

**Figure 4.**
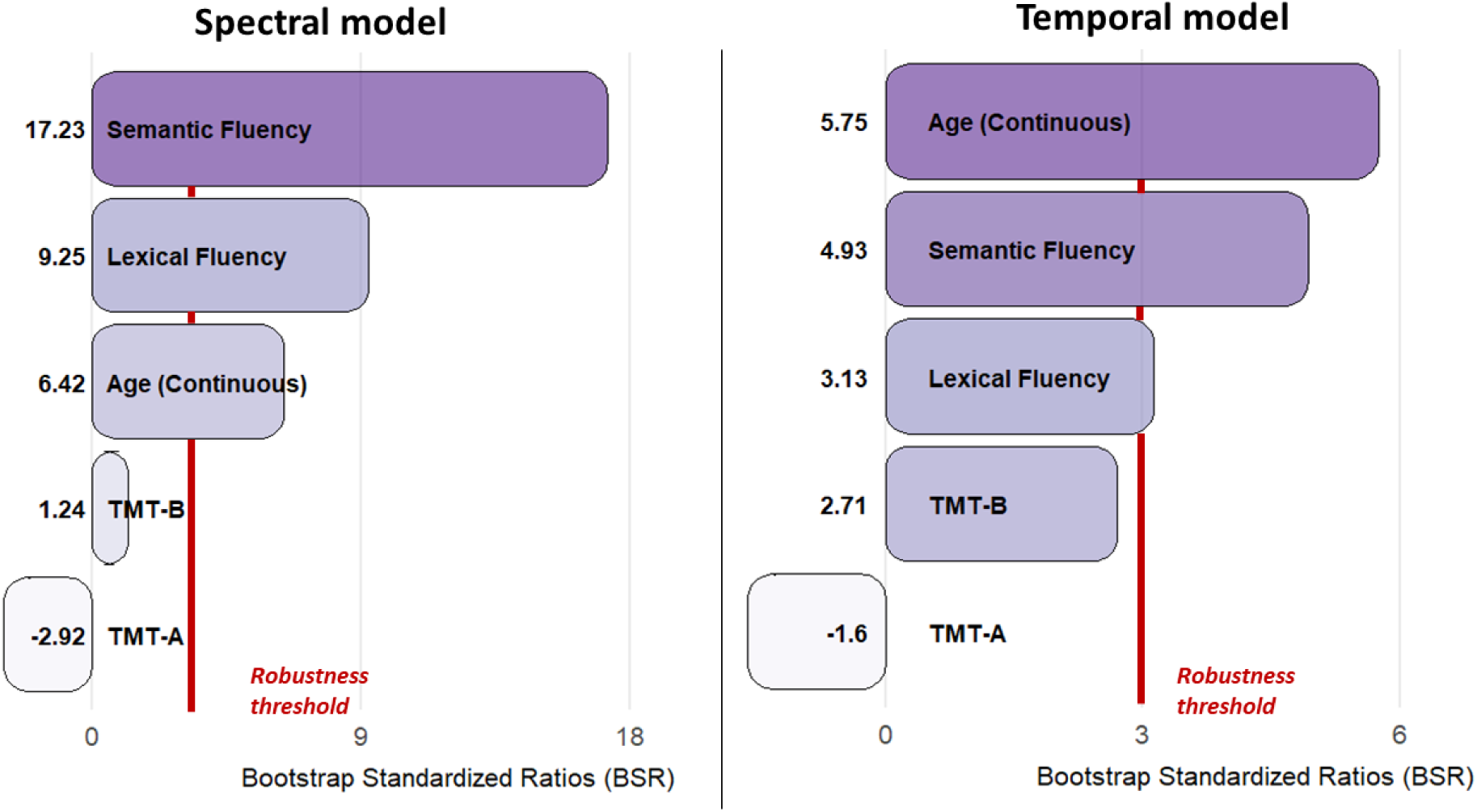
Salient demographic and behavioral features for each PLS model

Although not robust contributors (|BSR| < 3), we observed a positive trend for cognitive flexibility (TMT B), especially covarying with the brain state temporal organization (BSR*_temporal_* = 2.71). This lack of robustness could be attributed to high variability across subjects as evidenced by a high bootstrap standard deviation (0.2), twice to the average of other behavioral measures in the temporal model (∼ 0.1). By contrast, a negative trend was also observed for baseline processing speed (TMT A), especially covarying with spectral dynamics (BSR*_spectral_* = −2.92).

#### 3.2.2. Spectral model

Age and the associated behavioral profile (Fig. 4) covaried with a large-scale redistribution of oscillatory power across states (Fig. 5), particularly from the sensorimotor (State 2) and visuo-motor configurations (State 3) toward the states which bind with the higher-level DMN (State 1: semantic control & State 4: posterior right visual). This redistribution involved nearly the entire spectrum, although more robustly impacting the alpha (10-12 Hz), beta and low-gamma (20-37 Hz) bands. Additionally, we noted a less powerful desynchrony in State 5 (4 Hz peak & alpha 10-12 Hz band).

**Figure 5.**
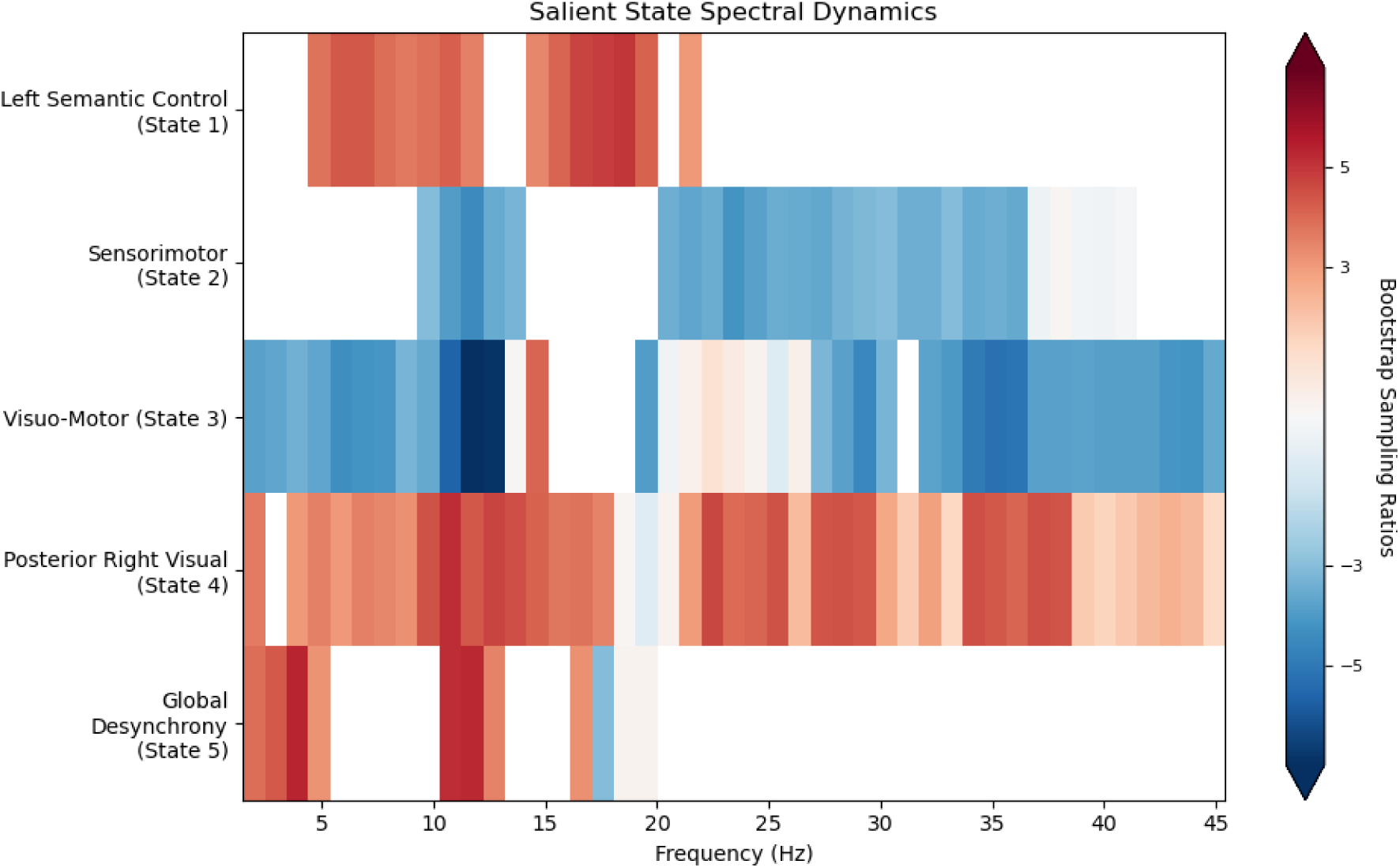
Salient spectral dynamics across brain states. A high BSR (red) indicates robust contribution to the latent brain variable, covarying with age and the behavioral profile shown in Fig. 4.

#### 3.2.3. Temporal model

The temporal PLS model provided complementary insights to the broad spectral changes identified previously, notably revealing changes in the processing stages of sentence generation (Fig. 6).

**Figure 6.**
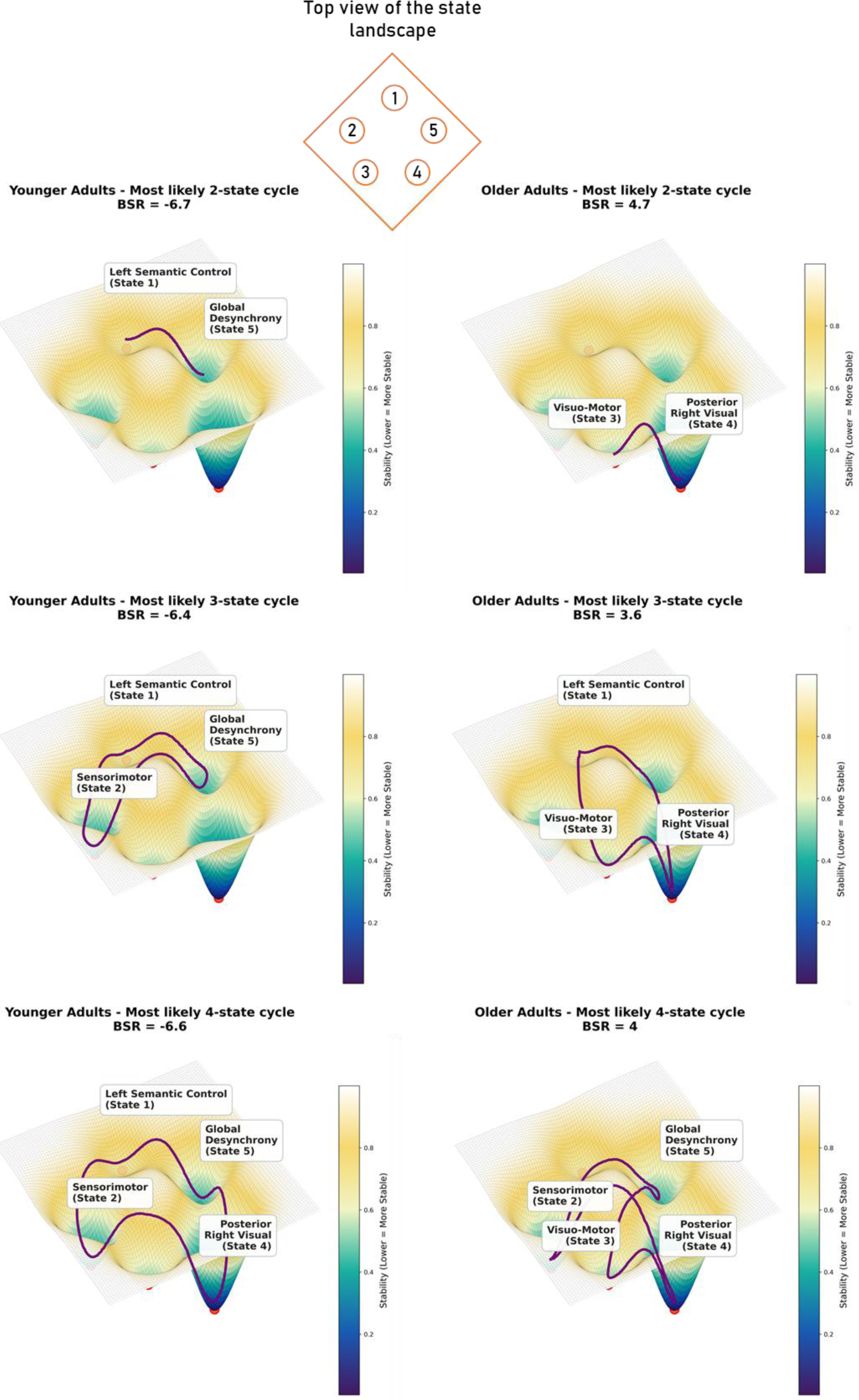
Most salient brain state cycles. Energy landscape. To visualize changes in cycle dynamics, we constructed a 3D energy landscape where states are organized in a circular layout (see Supplementary Information). The resulting landscape exhibits deep basins for states with high stability, and shallow valleys for more instable, transient states. **Projected cycle dynamics.** Salient brain state cycles are projected onto this landscape as smooth purple trajectories

While younger adults engaged in direct cycles involving the Left Semantic Control, Sensorimotor, and Desynchrony configurations (States 1, 2, 5), older adults exhibited a more segmented repertoire, additionally cycling through the Visuo-motor and Posterior Right Visual states (States 3 & 4). This segmentation in older adults suggests a form of temporal “chunking” of sentence generation stages, which covaried with better verbal fluency.

## 4. Discussion

This study investigated the neurophysiological mechanisms supporting naturalistic sentence generation in younger and older adults. Using Hidden Markov Modeling, we identified five co-active brain states during covert speech and demonstrated that their spatiotemporal properties are associated with age and verbal fluency. Taken together, our findings support our initial hypotheses. Healthy language aging appears to be characterized by an embodied semantic strategy, scaffolded by a coordinated time-frequency reorganization of large-scale brain dynamics spanning language-semantic, language-control, sensorimotor, and visual domains.

### Dynamical chunking: an age-related compensatory mechanism?

Our analysis of state-transition cycles complements recent evidence that large-scale brain networks organize into structured, nontrivial temporal sequences during cognitive tasks (Van Es et al., 2025). As illustrated in Figure 6, we observed a marked shift in cycle dynamics with age. Younger adults predominantly relied on a direct loop involving left-lateralized semantic control (State 1), sensorimotor processing (State 2), and global desynchronization (State 5) during sentence generation. Together, these states seem to support controlled word retrieval and articulatory-motor preparation. In contrast, older adults exhibited a segmented dynamic in which this loop was split into two sub-cycles, each interfacing with additional visuo-posterior configurations (States 3 & 4). These states appear to act as intermediary perceptual integration step between articulatory-motor processing (State 3: Visual-SMN) and higher-level semantic retrieval (State 4: Visual-DMN). This finding suggests that maintaining verbal fluency in older adulthood may require traversing a more fragmented functional trajectory that supports motor imagery (State 3) and visually guided lexical access (State 4) (Roos et al., 2023).

This contribution of visuo-posterior activity to sentence generation in aging provides novel task-based MEG evidence that older adults adopt an “embodied semantic strategy” for language production, in support of the SENECA model (Guichet, et al., 2026). It also aligns with the broader notion of predictive processing whereby multisensory inputs, here primarily visual, prime semantic representations to support complex cognition in aging (R. M. Brown et al., 2022; Ozkalp-Poincloux et al., 2026; Spreng & Turner, 2021).

From a metabolic standpoint, this predictive route can be interpreted as form of energy-efficient “temporal coding” that could emerge as “natural consequence of resource optimization” in older adulthood (Elias, 1955; Price & Gavornik, 2022). Since the aging brain faces a reduction in metabolic resources (Asimakidou et al., 2025), shifting from a streamlined three-state cycle to a segmented five-state dynamic may be a compensatory ‘chunking’ mechanism that reduces signaling costs during language production (Hechler et al., 2023). In this framework, our results suggest that visuo-posterior integration could act as an efficient mediator between distinct processing segments.

Although speculative, this interpretation aligns with evidence indicating that older adults primarily allocate metabolic resources to posterior hub regions (Deery et al., 2024). Prior resting-state MEG findings have also highlighted dorso-posterior brain states as candidate for information integration in age-related language production (Guichet et al., 2025).

#### Spectral mechanisms of the semantic strategy

The brain states’ oscillatory dynamics were also associated with the age-related semantic strategy. We primarily observed a dissociation between states primarily involved in sensorimotor-related processing (State 2: SMN; State 3: Visual-SMN) and those associated with semantic-related networks (State 1: semantic control; State 4: Visual-DMN) (Fig. 5). These state categories exhibited reciprocal spectral modulations with age: sensorimotor-related states exhibited broad suppression, whereas semantic configurations displayed robust power increases across frequencies.

One possible explanation is that, as the temporal sequence between semantic and sensorimotor processing becomes less efficient with age, the brain redistributes its resources to support increased engagement from DMN-semantic hubs. This interpretation is compatible with the LARA-C model (Baciu & Roger, 2024) which emphasizes the recruitment of compensatory semantic pathways to support language production in aging, and suggests that older adults resolve lexical competition at the semantic level to mitigate control deficits (Zheng & Piai, 2025). While such control deficits were not robustly identified (trending associations with TMT performance), we still observed reduced theta-alpha desynchronization in older adults across all major networks (State 5). This oscillatory dynamic aligns with recent evidence that alpha temporally coordinates theta processes (Beste et al., 2023), and could impact top-down control, attentional gating mechanisms (Foxe & Snyder, 2011; Pscherer et al., 2026), or perception-action integration (Eggert et al., 2025).

Nonetheless, it is important to note that the multivariate nature of PLS analysis limits the ability to isolate the independent contribution of specific frequency bands. Specifically, we did not find an expected association between beta power decreases and age-related semantic integration (Piai et al., 2014, 2015, 2020), but we observed enhanced alpha-band modulations in visuo-posterior states which could be interpreted as increased demands for semantic control in older adults (Guichet et al., 2025; Klimesch, 2012; Zioga et al., 2024). In sum, observed oscillatory changes suggest that age-related sentence production involves a coordinated, multi-spectral re-organization rather than the involvement of specific frequency bands.

### Limitations and Future directions

A primary limitation of this study is the relatively small sample size, particularly after exclusion of outliers in the older age group, which reduces statistical power and limits generalization. This constraint may partly explain the absence of more subtle effects with certain executive measures, such as TMT performance. In addition, the present covert sentence generation task may impose relatively low cognitive demands, which may have minimized larger age-group differences. Prior studies have shown that syntactic processing can remain preserved in older adults’ sentence production under low linguistic demands (Davidson et al., 2003; Hardy et al., 2022; Lee et al., 2022). Future research will be necessary to validate the proposed chunking mechanism in aging, by using more demanding linguistic paradigms that tap into working memory, such as complex lexico-syntactic constructions (Agmon et al., 2023; Sung et al., 2024), by eliciting sentences with abstract concepts to control for the degree of embodied semantic representation (Diveica et al., 2025; Pexman et al., 2023).

Although MEG provides excellent temporal resolution, signal-to-noise ratio limitations were evident, with the HMM framework isolating a low-frequency state likely reflecting noise. Moreover, future work could explicitly model for the aperiodic component of the signal, which is known to flatten with age (Mooraj et al., 2025). Beyond cortical dynamics, cortico-cerebellar loops and hippocampal contributions are increasingly recognized as central to predictive updating of internal models, especially during language processing (Casto et al., 2026; Yaghoubi et al., 2026). Integrating these subcortical sources, for example through MEG-fMRI approaches or invasive recordings such as stereo-EEG, will be essential to further elucidate the high-gamma band processing of embodied semantic representations in the aging brain (Dupont et al., 2025).

## Conclusion

This study investigated the MEG brain state dynamics supporting sentence generation in healthy aging using a cohort of 22 French participants. Overall, results suggest that semantic processing constitutes a compensatory adaptation for language production in aging, supported by a spatially distributed, spectrally diverse, and temporally segmented brain architecture.

This architecture is compatible with a ‘chunking’ mechanism that segments processing between lexico-semantic retrieval and articulatory-motor preparation in older adulthood. These processing stages may be interfaced by intermediary visuo-posterior activity, which could serve as a functional bridge within the processing sequence. Finally, we speculate that such an organization may contribute to a relatively resource-efficient functional layout in aging, helping to maintain language fluency despite a reduced metabolic budget.

## Data and code availability

Code is made publicly available at: https://github.com/LPNC-LANG/MEGAGING

## Supplementary information

### CRediT author statement

Conceptualization: MB, EC, SH; Investigation, Data Curation: VA, CL, SH, AC; Methodology, Formal Analysis & Software: CG, SH, RZ; Writing – Original Draft: CG; Writing – Review & Editing: All authors; Supervision & Funding Acquisition: MB

## Acknowledgments

We sincerely thank all participants for their contribution to the study. This work was supported by the Université Grenoble Alpes (IRGA Research Program) and the CDP CerCoG (ANR project ANR-15-IDEX-02).

