## Supplementary information for "MEG State Dynamics of Sentence Generation: Evidence for a Compensatory Chunking Mechanism in Healthy Aging"

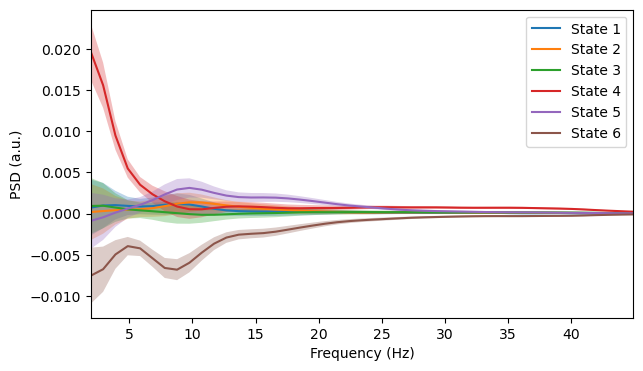
***Outlier Detection***

**Figure S1. Group-average inspection of the states’ PSD (before exclusion).** State 4 shows a low-frequency profile with high amplitude, suggesting aperiodic activity

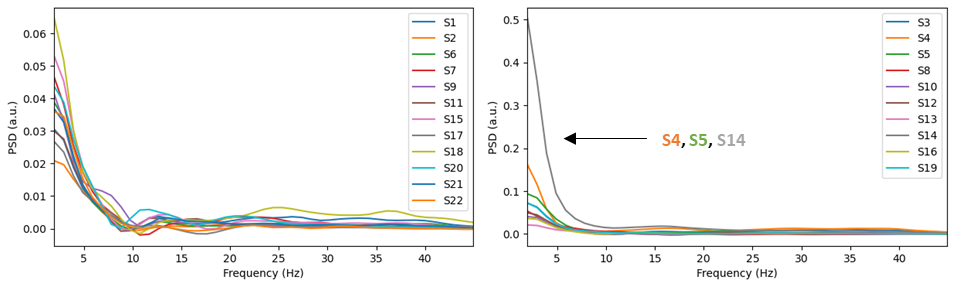

**Figure S2. Subject-specific inspection of State 4 (before exclusion).** Spectral profiles of younger subjects (left panel) and older subjects (right panel)

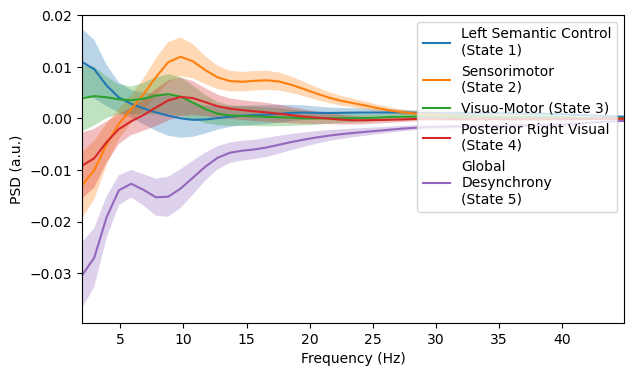

**Figure S3. Group-average inspection of the five states’ PSD.** We removed State 4 and refitted the GLM-spectrum to derive the PSD or each state. These PSDs correspond to the ones reported in Figure 2 in the main text, after excluding the three outlying subjects shown in Fig. S2 from the older age group: S4 (female, age 59), S5 (female, age 75), and S14 (female, age 60)

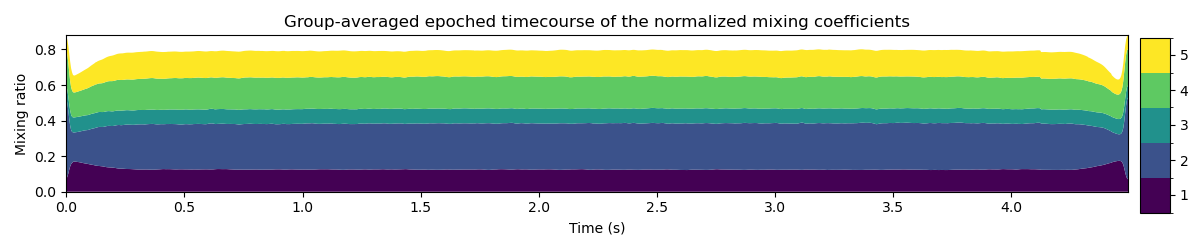

**Figure S4. Group-averaged epoched state time course**. This was obtained after epoching the states’ time courses across the 80 trials, and then averaging across subjects without the three outliers. The border effect is likely due to trial concatenation which created a discontinuity in the signal

***Visualizing cycle dynamics with a 3D energy landscape***

The landscape shown in Figure 6 was obtained by mapping the stationary distributions (i.e., the average time spent in each state) to a potential energy surface. Each of the five states was assigned a coordinate in a 2D plane using a circular layout. The energy at any point on the grid was defined as the negative weighted sum of the multivariate Gaussian kernels centered at these coordinates, where the weights are proportional to the state’s stationary probability. Salient trajectories from the PLS model were then projected using quadratic spline interpolation.
